# Decoding of EEG signals reveals non-uniformities in the neural geometry of colour

**DOI:** 10.1101/2021.06.17.448044

**Authors:** Tushar Chauhan, Ivana Jakovljev, Lindsay N. Thompson, Sophie M. Wuerger, Jasna Martinovic

## Abstract

The idea of colour opponency maintains that colour vision arises through the comparison of two chromatic mechanisms, red versus green (RG) and yellow versus blue (YB). The four unique hues, red, green, blue, and yellow, are assumed to appear at the null points of these the two chromatic systems. However, whether unique hues have a distinct signature that can be reliably discerned in neural activity is still an open question. Here we hypothesise that, if unique hues represent a tractable cortical state, they should elicit more robust activity compared to non-unique hues. We use a spatiotemporal decoding approach to reconstruct an activation space for a set of unique and intermediate hues across a range of luminance values. We show that electroencephalographic (EEG) responses carry robust information about isoluminant unique hues within a 100-300 ms window from stimulus onset. Decoding is possible in both passive and active viewing tasks, but is compromised when concurrent high luminance contrast is added to the colour signals. The efficiency of hue decoding is not entirely predicted by their mutual distance in a nominally uniform perceptual colour space. Instead, the encoding space shows pivotal non-uniformities which suggest that anisotropies in neurometric hue-spaces are likely to represent perceptual unique hues. Furthermore, the neural code for hue temporally coincides with the neural code for luminance contrast, thus explaining why potential neural correlates of unique hues have remained so elusive until now.

## Introduction

The idea of colour opponency maintains that colour vision arises through the comparison of two chromatic mechanisms, red versus green (RG) and blue versus yellow (BY). The four unique hues, red, green, blue, and yellow, are assumed to appear at the null points of these the two chromatic systems (Hering, 1920; Jameson and Hurvich, 1964; De Valois and De Valois, 1993). Colour vision starts in the retina, where light is absorbed in receptors (long-, medium, and short-wavelength sensitive cone receptors – L, M, S) and small bistratified ganglion cells that receive S-(M+L) cone input have been postulated to be the retinal origin of the BY channel, while midget ganglion cells that take the differences between the L and M cone output were believed to be the retinal origin of the RG channel (Lee et al., 2010). However, it has now been confirmed that the chromatic tuning of behaviourally characterised opponent channels differs from these early cone-opponent mechanisms, hence another transformation of chromatic signals must take place between the Lateral Geniculate Nucleus (LGN) and the primary or extrastriate visual cortex (De Valois and De Valois, 1993; Wuerger et al., 2005).

While some neuroimaging studies have attempted to identify a neural basis for unique hues, their results remain controversial. Stoughton and Conway (2008) reported neuronal clusters which were preferentially tuned to unique hues in the posterior inferior temporal (PIT) cortex of macaques. However, their findings have been challenged on the grounds that the study was not fully controlled for low-level differences in neuronal tuning, which could provide a more parsimonious explanation for their results (Conway and Stoughton, 2009; Mollon, 2009; Bohon et al., 2016). Similarly, Forder et al. (Forder et al., 2017a) reported that event-related potentials (ERPs) for unique hues show decreased latencies compared to non-unique hues. But the reported difference in peak latencies could, once again, have stemmed from differential activation of low-level, cone-opponent processes, to which ERPs are particularly sensitive (Rabin et al., 1994; Knoblauch et al., 1998). Thus, the neural basis of these cortical hue-opponent chromatic systems, and consequently, the unique hues, still remains an open problem.

One of the major reasons for the failure to address this issue has been the fact that neural activity is rich in coding possibilities which complicate our understanding of the relationship between external stimuli and the evoked response (Johnson, 2000; Jazayeri and Afraz, 2017). This is particularly true if potential low-level confounds can lead to a stronger, overlapping signal. This seems to be the case for unique hues, whose encoding is bound to overlap with, and be influenced by, the encoding of luminance contrast (see, e.g., Nunez et al., 2017). Ritchie et al. (2019) suggest that an ideal way to utilise neural decoding is to reconstruct an activation space from multivariate neural data and make psychological inferences by assessing whether such activation spaces correspond to psychological constructs. Recent studies have begun to apply this approach to challenges in colour neuroscience such as identifying the neural representations that underlie colour geometries (Rosenthal et al., 2021).

We hypothesise that, if there is indeed a distinct and discernible neural signature for unique hues, it should be reflected in the structure of the neurometric hue-representational space described by EEG signals. We used a decoding paradigm to test this hypothesis in two stages. First, we demonstrate that under isoluminant conditions, hue information can indeed be extracted from EEG signals, and that crucially, the encoding for unique hues is more robust than non-unique hues. To establish that our predictions generalise beyond a single decoding context (stimulus or task-wise), we test our decoding prediction using both active and passive viewing tasks. Second, we show that the structure of the neurometric space which encodes hue is distorted in the local neighbourhood of unique hues – suggesting a correspondence between low neural variability and unique hue percepts. Taken together, our findings suggest that the neural basis of perceptual unique hues is likely to be a set of stable fixed-points of a spatiotemporal population code for colour representations in the cortex.

## Methods

### Participants

In Experiment 1, twenty participants (16 females, 4 males) completed the study, ranging in age from 18-38 y.o. a. (mean age 21 y.o.a). In Experiment 2, 16 participants (all female) completed the study, ranging in age 19 – 32 years old (mean age 22 years). All participants reported normal or corrected-to-normal visual acuity, In Experiment 1, their colour vision was verified using the Trivector Cambridge Colour Test (Regan et al., 1994). In Experiment 2, we relied on the City University Colour Vision Test (Fletcher, 1975). Participants gave written informed consent and were reimbursed for their effort and time. The study was approved by the ethics committee of the School of Psychology, University of Aberdeen, and was in accordance with the Declaration of Helsinki.

### Stimuli and Procedure

The experiments were programmed using the CRS Toolbox and Color Toolbox (CRS, UK) for MATLAB (Mathworks, USA). In Experiment 1, stimuli were rendered on a 21-inch Viewsonic P227F CRT Monitor which was placed 70 cm away from the participant. The monitor was controlled through a Visage system (CRS, UK) and calibrated using ColorCAL2 (CRS, UK). Colours were generated on the basis of measurements taken with a SpectroCAL (CRS, UK). Participants gave their responses using a Cedrus R530 response box (San Pedro, USA). In Experiment 2, colours were presented on a Display ++ (CRS, UK) device. Responses were recorded using a CT-6 button box (CRS, UK).

Different sets of colours were used in the two experiments. In Experiment 1, stimulus colours were selected from a normative dataset (Wuerger and Xiao, 2015) (see Supplementary **Figure S3A**). The hue angles for unique red (UR) and unique green (UG) stimuli corresponded to mean values in the dataset – in the perceptually uniform CIE 1976 UCS space (Schanda, 2016), hue angles for the UR and UG stimuli were 14.4° and 133.4° respectively. Orange and turquoise stimuli were chosen such that orange (hue angle 41.5°) was the intermediate hue between UR and unique yellow, and turquoise (hue angle 185.1°) was the intermediate between UG and unique blue. All four stimuli were equally saturated in the CIE 1976 UCS plane. Three stimulus luminance levels were used: nominal iso-luminance (24 cd/m^2^), 45% Weber contrast (34.8 cd/m^2^) and 90% Weber contrast (45.6 cd/m^2^). In Experiement 2, stimuli consisted of participants’ subjective settings for two unique hues (yellow and green) and two intermediate hues (orange and turquoise), and hues situated 10° to the left and right of the subjective settings (in CIELCh colour space). Thus, for each observer, we effectively had four clusters of colours corresponding to the hues orange, yellow, green, and turquoise, with each cluster consisting of the observer setting for that hue, along with two flanking colours ±10° from the setting (e.g., unique yellow, a yellow 10° counter-clockwise and a yellow 10° clockwise). All colours were nominally isoluminant with the background (CIE 1931 xyY coordinates: 0.3127, 0.3290, 22.93 *cd*/*m*^2^).

EEG data was recorded during a shape discrimination task. The purpose of the task was to engage participants’ attention in a stimulus dimension orthogonal to colour - i.e., shape. The stimuli consisted of uniformly coloured shapes shown against a grey background. Each trial began with the appearance of a fixation cross, followed by a 2-degree circular stimulus (passive viewing event) which changed shape (shape change event) into either a diamond or a square (**Figure 1**). The passive viewing event occurred 700±200 ms after the appearance of the fixation cross, and the shape change event occurred 800-1500 ms after the passive viewing event.

**Figure 1:**
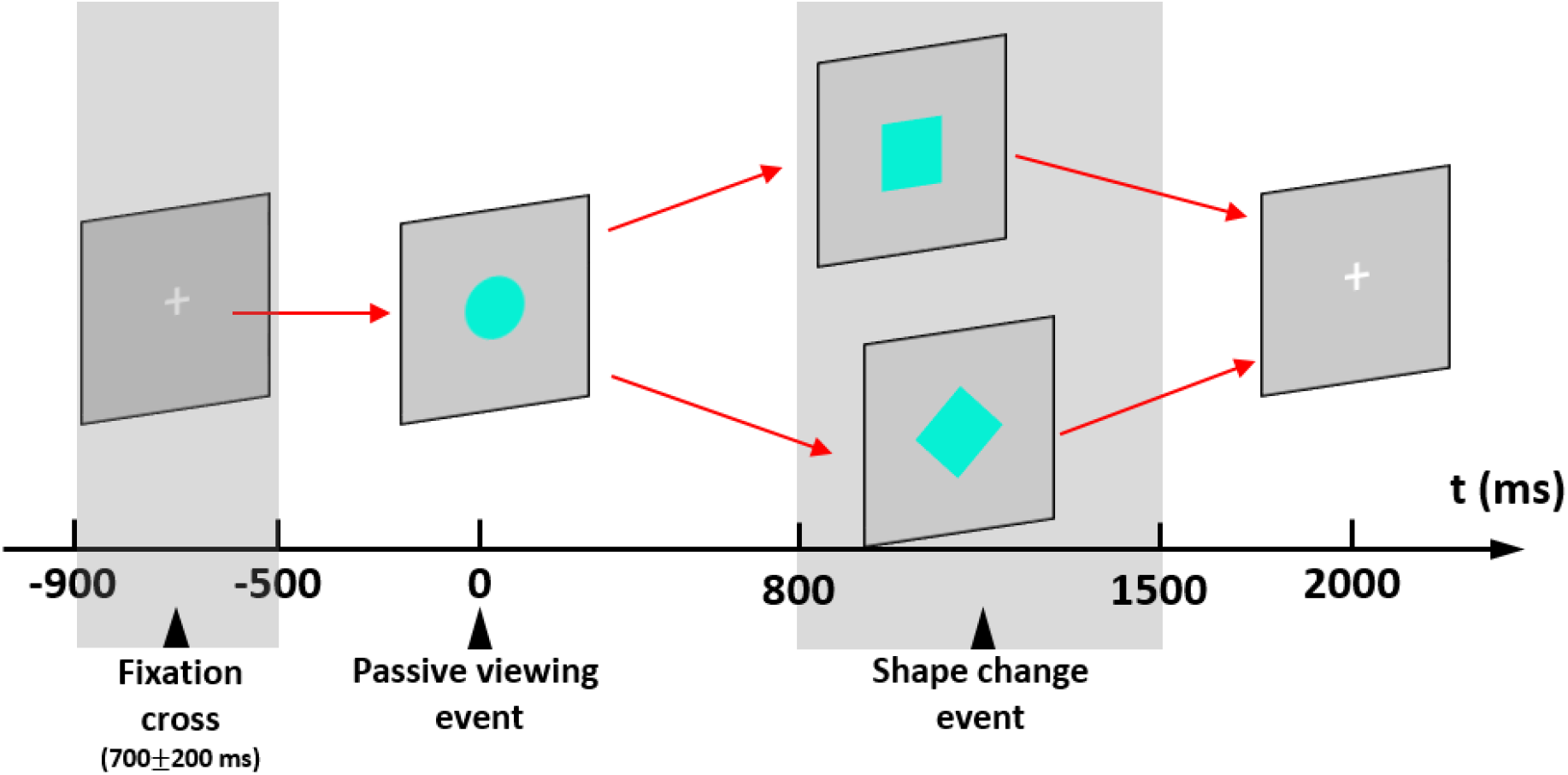
Experimental design. Each trial started with the appearance of a fixation cross, which was followed by the presentation of a circular uniformly-coloured stimulus at a random offset of 700±200 ms. At a random time-point 800-1500 ms after stimulus onset, the shape of the stimulus changed from circular to either square or diamond. Participants were instructed to discriminate the final shape via a button press as quickly and as accurately as they could. Each trial ended 2 seconds after stimulus onset. Two events were defined during each trial: a passive viewing event defined by the appearance of the stimulus, and a shape change event defined by the change in stimulus shape.

Participants identified the final shape of the stimulus using the left or the right button on a button box. The assignment of button to the target shape was counterbalanced between participants. The conditions were randomly intermixed, with a different order for each participant. The entire experiment was conducted in a sound-attenuated, electrically shielded chamber, with the screen being the only source of light. In addition to EEG recordings (described below), two other task-related variables were measured – task accuracy and reaction time. For each colour and shape combination, we had 30 trials. As diamond and square shape-change trials were subsequently collapsed together, this resulted in 60 trials per colour and 720 trials in total, presented in random order and divided into 10 blocks. This was the same for both experiments. In addition, Experiment 1 was preceded by a practice of 24 trials, while Experiment 2 was preceded by a practice of 16 trials. The EEG task took approximately 50 minutes to complete.

After the completion of the EEG experiment, participants rated each colour on a 9-point Likert scale for its representativeness of its category. This took approximately 5 minutes. Participants were asked to imagine the perfect representative for a colour category and rate how representative a sample was of that category, with 1 being the least representative and 9 being the most representative. All colours were displayed simultaneously on the screen during this procedure and remained on the screen until the participants completed the task. Colours were presented on the computer screen as a set of 4 rows of squares that showed the three luminance (Experiment 1) or hue (Experiement 2) values for that colour.

There were also two additional measures, specific to each experiment. In Experiment 1, for each participant, heterochromatic flicker photometry (HCFP) at 20 Hz (Walsh, 1958) was used to establish the departure from isoluminance for all colours. The task required the participant to adjust the luminance of the colour until perceived flicker was minimised. Participants performed 8 trials per colour – the step size was 0.5 cd/m2 and the flicker started from a randomly determined point that could be five steps above or below nominal isoluminance. These measurements were conducted to evaluate any individual differences in the amount of luminance contrast effectively present in nominally isoluminant stimuli. Rabin et al. (1994) demonstrate that departures from isoluminance need to be substantial to influence chromatic visual evoked potentials. Collecting HCFP data enabled us to verify that small departures from effective luminance did not significantly influence the efficiency of colour decoding.

Experiment 2 began with a hue adjustment task, in which participants made their individual hue settings for two unique hues (yellow and green) and two intermediate, non-unique hues (orange and turquoise). Participants performed one block of eight trials for each hue. The order of blocks (yellow, green, orange, turquoise) was randomized for each participant. Colours were defined in CIE LCh colour space to have the same chroma (C=25) and lightness (L=55). Initial hue angles were randomised to the following values: 90°+/− 12° for yellow, 180°+/− 12° for green, 45°+/− 12° for orange and 225°+/− 12° for turquoise. A coloured 2° circle was shown in the middle of the computer screen. Participants used the right and left buttons to change the hue along the CIE LCh hue circle in steps of 2° clockwise and counter-clockwise, respectively. Once the participants were happy with their setting, they completed the adjustment by pressing the top button. The task took approximately 10 minutes to complete. The first 6 participants performed the task without context. For the following 13 participants, we also presented a colour palette consisting of 19 squares 1° in size that ranged +/− 45° around the initial hue value, in steps of 5° of hue angle, positioned at 8.22° above the central stimulus. The colour palette provided context for the hue setting task. A between-subject ANOVA showed no difference in unique hue settings with and without context (F(1,14) = 0.23; p = .64, ηp^2^ = .02).

In total, the experiments lasted between two and a half and three hours, including the time to set up and remove the EEG electrodes.

### EEG recording and pre-processing

Continuous brain activity was recorded from 64 scalp locations using active Ag-AgCl electrodes and 4 ocular channels (providing VEOG and HEOG) connected to a BioSemi Active-Two amplifier system (BioSemi, Amsterdam, The Netherlands) at a sampling rate of 256 Hz. Data processing was performed using EEGLAB (Delorme and Makeig, 2004) for Matlab (Mathworks, UK). Epochs lasting 900 ms were extracted: 200 ms before the relevant event (stimulus onset or shape change) and 700 ms afterwards. Data was low pass filtered at 40 Hz. All trials with incorrect answers were excluded prior to the analysis. Artifact removal was then performed by using the FASTER toolbox (Nolan et al., 2010), the ADJUST toolbox (Mognon et al., 2011), and self-written procedures in MATLAB. FASTER is an automated procedure that detects contaminated trials and noisy channels that need interpolation (either in the entire EEG recording or on any single trials) by calculating statistical parameters of the data and using a *Z* score of ±3 as the metric that defined contaminated data. ADJUST is an automated procedure that operates on maps resulting from independent component analysis of EEG data, using properties of these components to label them as eye blinks, vertical or horizontal eye movements, or channel discontinuities so that they can be subtracted from the recording. We first rejected trials with global artifacts using FASTER, then ran an independent component analysis and applied ADJUST to the obtained decompositions, and finally, conducted channel interpolation with FASTER. In addition, any trials with eye movements were rejected based on ±25μV deviations from the horizontal electrooculogram in the uncorrected data. Blinks were rejected using a thresholding procedure similar to FASTER (Junghöfer et al., 2000).

Incorrect and rejected trials amounted to a very small proportion of the data – in Experiment 1, between 1% and 13% of total trials, and in Experiment 2, between 3% and 17% of total trials.

### EEG classification

The classification of EEG signals was set up as a set of time-windowed error-correcting output codes models (tECOC) operating on 20ms snippets of the signals (other reasonable time-windows yielded similar results, see Supplementary **Figure S1A**) from the occipital electrodes (the entire set of 64 electrodes yielded similar results, see Supplementary **Figure S1B**). Linear discriminant analysis (LDA) classifiers were employed as learning units due to their relative simplicity and computational efficiency. Denoting the EEG activity as a random multivariate variable ***X***, and the stimulus label (colour and/or luminance) by the random variable *Y* (where realisations of *Y* are drawn from the set of all possible labels denoted *L*), the probability that the observed activity ***x*** is elicited by the stimulus described by label *y* is given by the Bayes rule:

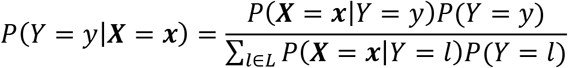

In LDA, the likelihood term is estimated by a multivariate Gaussian density function:

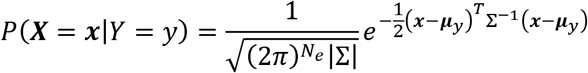

Here, *N*_*e*_ is the number of electrodes, ***μ***_*y*_ is the mean EEG activity for the label *y*, and Σ is the covariance matrix of the activity. The log-posterior objective function *δ*_*y*_(***x***) for the label *y* can thus be written as:

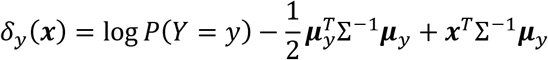

Data for each observer was modelled separately, and the whole process was repeated 10 times. In each repetition for any given observer, the data were split into 5 folds containing roughly equal number of samples for each label. Each of the five folds was then tested by training the model on the remaining 4 folds.

tECOC analysis gave us a time-series of confusion matrices (CMs) which characterise the model performance over the duration of the trial (see Supplementary **Video V1**). At each time-point, while the diagonal of the CM gives a measure of model accuracy (true positive rate), the off-diagonal elements represent misclassifications, which are crucial towards understanding the topography and information content of the representational space (see Representational Similarity Analysis below). In addition, a permuted model was also trained using a shuffled set of labels to estimate empirical chance performance. The empirical chance performance was found to be close to theoretical chance level under the assumption of equilikelihood (see Supplementary **Figure S1C**). The statistics (both within-observer and population) were calculated at each time-point by comparing model performance with empirical chance performance of the ‘permuted’ models using a two-tailed randomisation test with 1000 permutations.

### Representational Similarity Analysis

The time-series of confusion matrices estimated by tECOC models were used to calculate pairwise dissimilarities between stimulus classes. Given a confusion matrix *C*, where each element *c*_*ij*_ denotes the probability of the stimulus type *i* being labelled as *j* by the model, first, a label-normalised matrix *S* was constructed such that *s*_*ij*_ = *c*_*ij*_/*c*_*ii*_. This asymmetric measure was then used to calculate a symmetric dissimilarity tensor Δ_tECOC_ given by

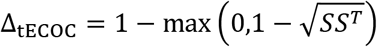

Here, the geometric mean across stimulus pairs is used to generalise distances in representational space (Shepard, 1958; Kaneshiro et al., 2015). A similar estimation was also made for the EEG data using a more traditional dissimilarity metric given by

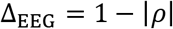

Here, *ρ* is the Pearson-correlation matrix for the EEG responses elicited by the stimuli. Finally, pairwise differences in CIELAB hue angles of the stimuli were used to estimate a perceptual dissimilarity matrix. The perceptual dissimilarity was compared to Δ_*tECOC*_ and Δ_*EEG*_ using rank-correlation estimates (Kendall’s coefficient).

## Results

### Experiment 1: Decoding unique and intermediate hues with and without luminance contrast

We measured EEG signals in a cohort of 20 participants while they viewed coloured stimuli (coloured shapes on a grey background) consisting of two unique hues – unique green and unique red, and two non-unique hues – orange and turquoise. In each trial, a coloured disc changed shape to a diamond or a square at a random time-point 800-1500ms after stimulus onset (**Figure 1**). The participant’s task was to identify the target shape. The stimuli were either isoluminant with the background (0% luminance contrast), or presented at 45% or 90% luminance-contrast. This gave us a dataset of EEG signals labelled both in hue and luminance-contrast. The task was easy, resulting in high overall accuracy (95% ± 1% SE, see Supplementary **Figure S4A**) and very fast responses (mean response time (RT) of 462 ± 15 ms, see Supplementary **Figure S4B**). Response-time data was analysed with a 3×4 repeated measures ANOVA (3 levels of luminance contrast vs. 4 hues), which yielded a significant main effect of luminance contrast (*F*(1.49, 28.28) = 67.56, *p* < 0.001, *μp*^2^ = 0.78) and interaction with hue (*F*(6, 114) = 3.56, *p* = 0.003, *μp*^2^ = 0.16) – hue itself did not have an effect (*F*(1.88, 35.79) = 2.93, *p* = 0.07). We deconstructed the interaction by performing separate repeated measures ANOVAs at each luminance contrast: while at isoluminance there was a significant effect (*F*(3, 57) = 6.19, *p* = 0.001, *μp*^2^ = 0.25) driven by slower RTs for green (vs. red *P* = 0.019; vs orange *P* = 0.008, vs. turquoise *P* = 0.003), there were no differences at 45% luminance contrast (*p* = 0.16) or at 90% luminance contrast (*p* = 0.11).

After the completion of the EEG experiment, participants rated each colour on a 9-point Likert scale for its representativeness of its category (red, orange, green or turquoise). The average ratings and their SEs were as follows (see Supplementary **Figure S4C**): isoluminant red 4.35 ± 0.48; red at 45% luminance 2.85 ± 0.32; red at 90% luminance 1.90 ± 0.23; isoluminant green 7.70 ± 0.23; green at 45% luminance 6.10 ± 0.35; green at 90% luminance 5.55 ± 0.43; isoluminant orange 3.75 ± 0.48; orange at 45% luminance 4.15 ± 0.43; orange at 90% luminance 3.60 ± 0.32; isoluminant turquoise 6.00 ± 0.47; turquoise at 45% luminance 6.75 ± 0.38; turquoise at 90% luminance 6.40 ± 0.5.

#### Unique hues can be robustly decoded from EEG signals

First, we asked whether the measured EEG waveforms contain consistent, discernible information about the hue of the stimulus. To do this, we trained tECOC models for each observer using only EEG responses to isoluminant stimuli, as this ensured minimal interference from luminance-contrast signals. In the first instance, we performed this analysis for epochs defined by the passive viewing event. We found that within a 100-300 ms window after stimulus onset, both unique hues could be decoded with above-chance accuracy (**Figure 2A**). The non-unique hues, on the other hand, showed a much lower score (**Figure 2B**). This pattern is stable over a range of tECOC time-windows (Supplementary **Figure S1A**) and also holds when the entire set of 64 electrodes is used (Supplementary **Figure S1B**). The presence of signal on all electrodes is not surprising – unlike functional magnetic resonance imaging (fMRI), EEG does not detect localised physiological activity in a volume, but instead picks up a linear superposition of signals from a range of physiological sources. Thus, the signal is present in some degree at all sensors, with its amplitude (and thus also its signal to noise ratio) dependent on the position of the sensor relative to the source(s) (see, e.g., Maris, 2012 for a discussion of the so-called common pick-up problem).

**Figure 2:**
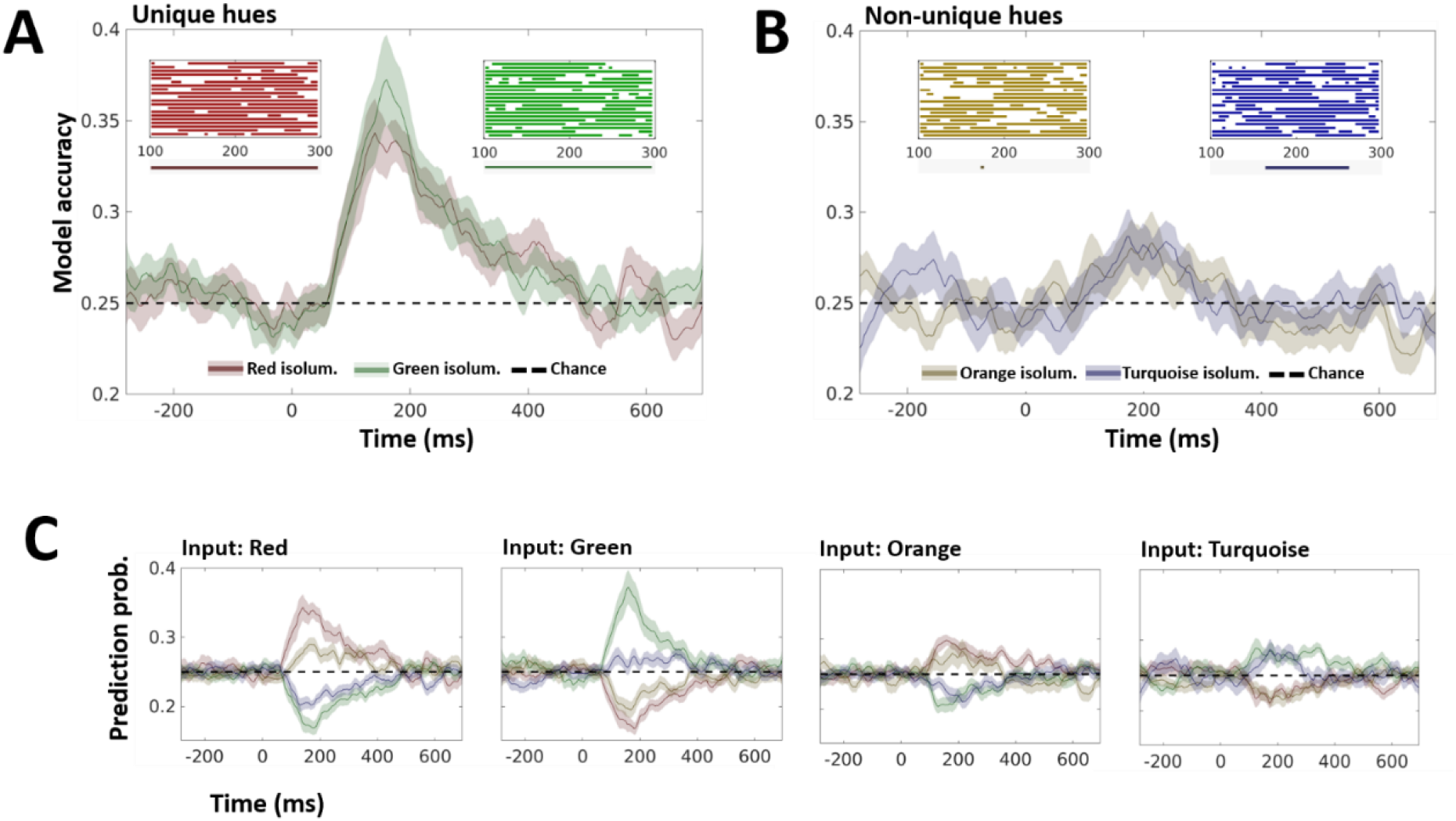
Decoding isoluminant Unique and Non-unique hues from EEG responses. A tECOC classification model was trained on EEG responses recorded in *N* = 20 participants as they viewed isoluminant stimuli (Unique Hues: red and green; Non-unique hues: orange and turquoise). **A.** Model accuracy for Unique hues. This corresponds to presenting the trained model with EEG responses to Unique Hue stimuli and estimating the probability with which the model is able to determine the correct stimulus hue (diagonal of the confusion matrix). The two solid lines show the mean accuracy of the model at each time-point. The hues are colour-coded, with the red and green lines representing model accuracy for unique red and unique green stimuli respectively. The shaded regions around the lines show ±1 standard-error of the mean over the 20 observers. A dashed line indicates the theoretical chance performance of the model (the empirical chance performance closely followed the theoretical chance level, and is shown in Supplementary **Figure S1C**). The two inlays show the classification accuracy (top-left: unique red, top-right: unique green) of models trained for each of the 20 observers. Only 100-300 ms after stimulus onset are shown in the inlays. The solid lines show the period when the classification performance was significantly above chance (*p* < 0.05 in a permutation test comparing observer-model performance to the performance of a model trained on randomly shuffled labels). Under the x-axis of the inlays, a solid line shows when the mean accuracy calculated across all observers was above chance (*p* < 0.05 in a permutation test comparing average performance over all observers to the average performance of models trained on random labels). **B.** Model accuracy for non-unique hues. The accuracy of the model for non-unique hues is shown in a manner analogous to **A**, with the orange and blue colours representing the orange and turquoise stimuli respectively. **C.** Misclassification probabilities. Given the EEG response (at a given time-point) to one of the four hues, the model can either make an accurate prediction of the label (panels **A** and **B**), or misclassify the input. Each of the four subpanels here shows the prediction probabilities for one particular input label (shown on the top-left, above each subpanel), thus corresponding to one row of the confusion matrix. For instance, the first subpanel shows the probabilities (at each time-point) that the model classifies EEG responses to unique green stimuli as being elicited by unique green (accuracy), unique red, orange or turquoise stimuli. The colour coding for the four stimulus hues in each subpanel is the same as panels **A** and **B**. Also see Supplementary **Video V1**, which shows how the confusion matrix changes as a function of time elapsed from stimulus onset.

For each participant, we also measured subjective isoluminance for each stimulus colour (see *Methods* for details). While one participant did not understand the task, the means, SEs and ranges of the settings from the remaining 19 participants were as follows: red 0.14 ± 0.57 *cd*/*m*^2^ (−6 to 5.25 *cd*/*m*^2^); green −1.09 ± 0.49 *cd*/*m*^2^ (−6.58 to 1 *cd*/*m*^2^); orange 0.08 ± 0.56 (−4.34 to 6.50 *cd*/*m*^2^); turquoise (−0.05 ± 0.65 *cd*/*m*^2^ (−7.08 to 7.83 *cd*/*m*^2^).

Model accuracy quantifies the ability of the model to correctly identify the hue of a stimulus when presented with the corresponding EEG response. Theoretically, it is the sum of hit rates (true positive rates) for all labels, and corresponds to the diagonal of the confusion matrix. However, a deeper insight into model performance can be obtained when, in addition to the detection accuracy for a given input class, one also considers the probability of misclassification of inputs from this class. To investigate this, we estimated the off-diagonal elements of the confusion matrix. This allowed us to infer which classes are most likely to be confused by the model – thus providing a means of understanding how similar the information contained in EEG signals corresponding to different hues is. The subpanels of **Figure 2C** (see also Supplementary Video V1) show the probability (over time) with which the model assigns each of the four hue labels to EEG responses elicited by a given input hue (the input hue is labelled above each subpanel). Thus, each subpanel in **Figure 2C** shows one row of the confusion matrix. Within a 100-300 ms window, each input hue is only confused with its proximal pair (red and orange, and green and turquoise), while the prediction probabilities for non-proximal hues are below chance. This is also reflected in the checkerboard-like pattern observed in Supplementary Video V1. Furthermore, the model is likely to label EEG responses to non-unique hues (orange and turquoise) as being elicited by their proximal unique hues (red and green respectively) with almost equal probability, but not vice-versa. Once again, this suggests that EEG signals between 100-300 ms carry more robust representations of unique hues compared to non-unique hues.

The passive viewing at trial outset was followed by a change in the shape of the stimulus from a circle to either a square or a diamond at a random time-point 800-1500 ms from stimulus onset (see **Figure 1**). The colour of the stimulus was task-irrelevant, and the hypothesis here was that since the observer will be attending to the stimulus shape, the EEG signal would be qualitatively different between the passive and shape-change segments. This would, in-turn, allow us to test if this difference is reflected in the ability of the model to classify hue information in the signal. It has been argued that colour-related activations should still be observed as long as the hue remains unattended and task-irrelevant (Forder et al., 2017b). To test this hypothesis, we trained tECOC models on the epochs defined by the shape-change event. As expected, the two segments were found to elicit activity which differed significantly both in the sequence of ERP peaks as well as topography (**Figure 3A**). However, despite this difference, we were able to perform hue detection during the shape-change segment with an accuracy very similar to the passive viewing segment – both in terms of peak decoding score and its time-course (**Figure 3B**). This suggests that the temporal structure of the hue-related information in EEG signals is indeed robust to changes in the task (as long as the hue itself remains task-irrelevant), and can be extracted even when the observer is engaged in a concurrent shape discrimination task.

**Figure 3:**
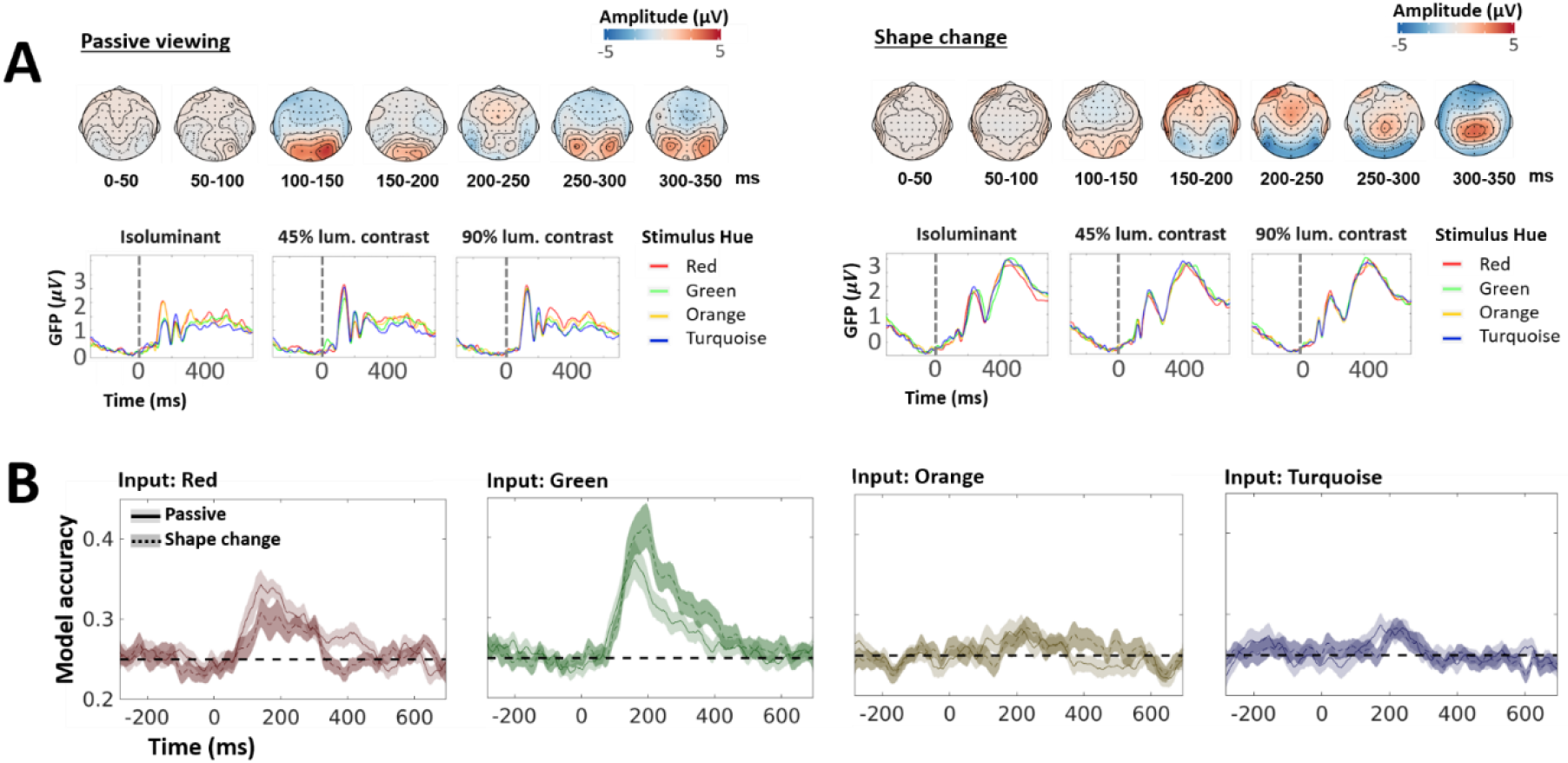
Decoding performance for active and passive tasks is very similar, despite large differences in stimulus-evoked activity. **A.** Global field power (GFP). The left side of the panel shows the topographies and the GFPs for stimulus onset. The hues are colour-coded (unique green is shown in green, red in red, etc.), and each panel shows the GFP for one luminance-contrast condition. The stimulus onset is marked by a dashed line at 0 abscissa. The right side of the panel shows the same for the shape-change event. **B.** Robustness to task. Separate models were trained using passive viewing and shape-change segments. Each subpanel shows the accuracy of the two models for one particular input hue (e.g., the leftmost panel shows the model accuracy when EEG responses to red stimuli were used as inputs). The performance of the passive-segment model is shown using the same colours and symbols as Figure 2A, while the shape-change model is shown using a dashed line for observer mean and darker shading for ±1 standard-error of the mean.

#### Luminance signals interfere with chromatic information in occipital ERPs

Next, we investigated whether hue identity could still be decoded when both chromatic and luminance information was present in the EEG signal. A chromatic-driven ERP is characterised by a robust negative deflection at about 120-220 ms after stimulus onset (Murray et al., 1987; Berninger et al., 1989; Tobimatsu et al., 1996), but this response is significantly altered by the addition of luminance contrast (Rabin et al., 1994). According to normative work by Rabin et al. (1994), while observer isoluminance drives ERPs in a manner closely resembling nominal isoluminance, any substantial changes in luminance contrast have been found to result in highly dissimilar waveforms. To assess if this would also impact classifier performance, we constructed a model that evaluated how decoding performance was affected when the model was trained on inputs which differ not only in hue but also luminance-contrast. We trained tECOC classifiers for each observer using 12 labels, corresponding to three different luminance-contrast levels for each of the four hues. In **Figure 4**, we present the performance of our model in a manner similar to **Figure 2C**. Each panel is one row of the confusion matrix, i.e., given the EEG signals for an input stimulus, it shows the prediction probabilities for all 12 labels. The hue of the input is denoted by the row (labelled in the right margin) and its luminance-contrast by the column (labelled on top). The same colours as **Figure 2C** are used to denote the four hues. In addition, for each hue, we also use two additional brightness levels to represent the two luminance contrast ratios (thus, for a given hue, isoluminant stimulus is the least bright, 45% luminance contrast is brighter, and 90% luminance contrast is the brightest). We find that while isoluminant signals can indeed be classified 100-300 ms after stimulus onset (left column), addition of luminance information disrupts the model performance for all hues (middle and right columns). Furthermore, we find that the classifier does not confuse isoluminant and non-isoluminant stimuli. This suggests that in contrast to a change in stimulus-shape where the temporal structure of hue-related information was preserved, addition of luminance-contrast to the stimulus disrupts the temporal patterns which encode hue-information.

**Figure 4:**
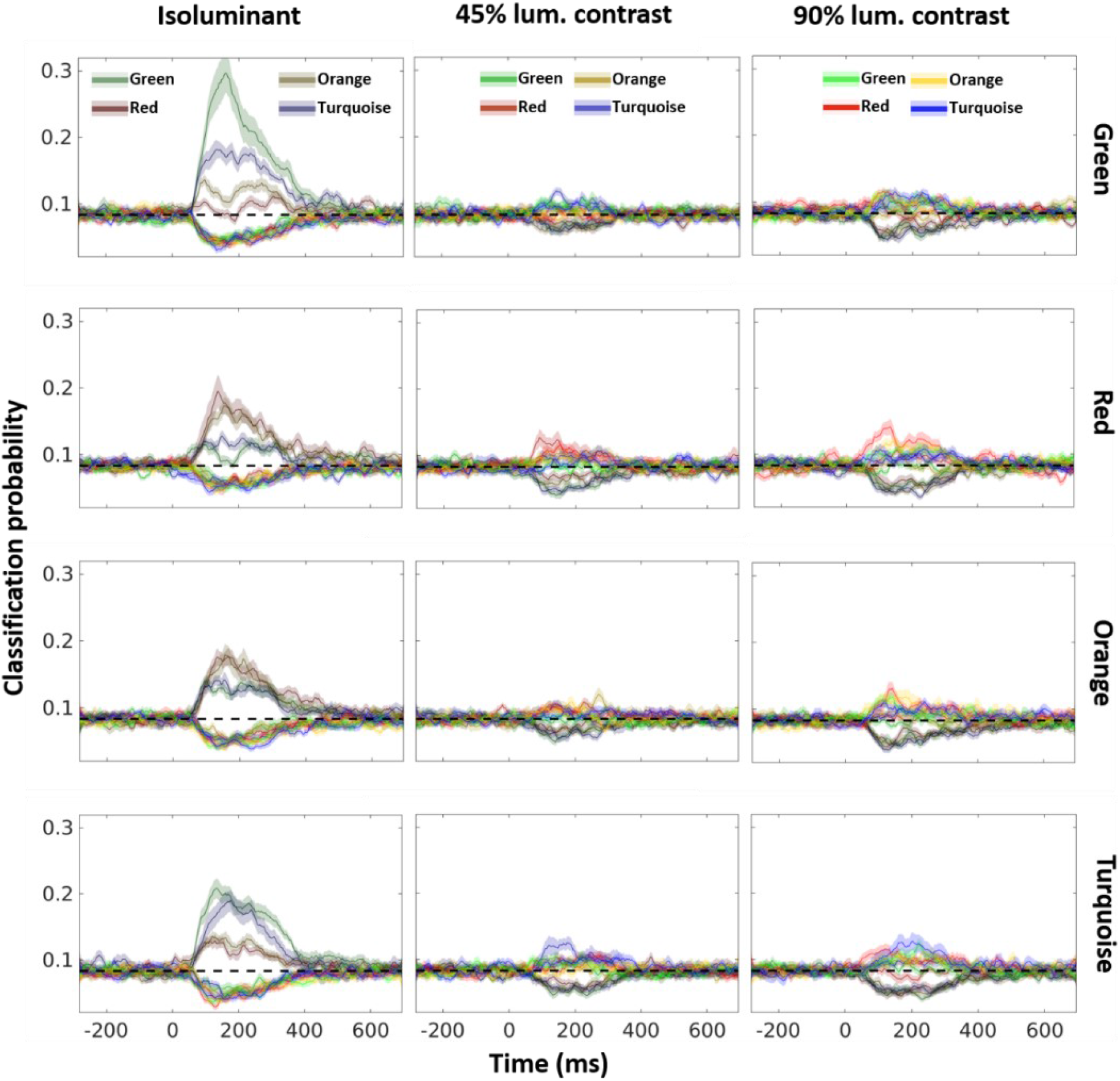
Luminance information disrupts hue decoding. EEG responses to unique (green and red) and non-unique (orange and turquoise) hues at three luminance-contrasts (isoluminant, 45% and 90%) were used to train a tECOC model. Each of the 12 subpanels in this figure represents one row of the confusion matrix (similar to Figure 2C). This corresponds to presenting the trained model with EEG responses to a given stimulus class, and observing the classification probabilities for all classes, including the input class. The hue and luminance contrast of the input labels are denoted by the row and column respectively. For each predicted label, the hue is represented by the corresponding colour (green, red, orange and turquoise), and the luminance-contrast by the brightness (isoluminant: lowest brightness, 45% contrast: intermediate brightness, 90% contrast: highest brightness).

To characterise the effect of luminance, we trained a model using only the luminance labels of EEG signals (i.e., we used three labels corresponding to the three contrast levels). We found that all luminance conditions (**Figure 5A**) can be decoded to above-chance levels. An examination of the misclassification patterns of the model (**Figure 5B**) further revealed that while isoluminant stimuli are robustly classified, the non-isoluminant conditions are more likely to be confounded with one another.

**Figure 5:**
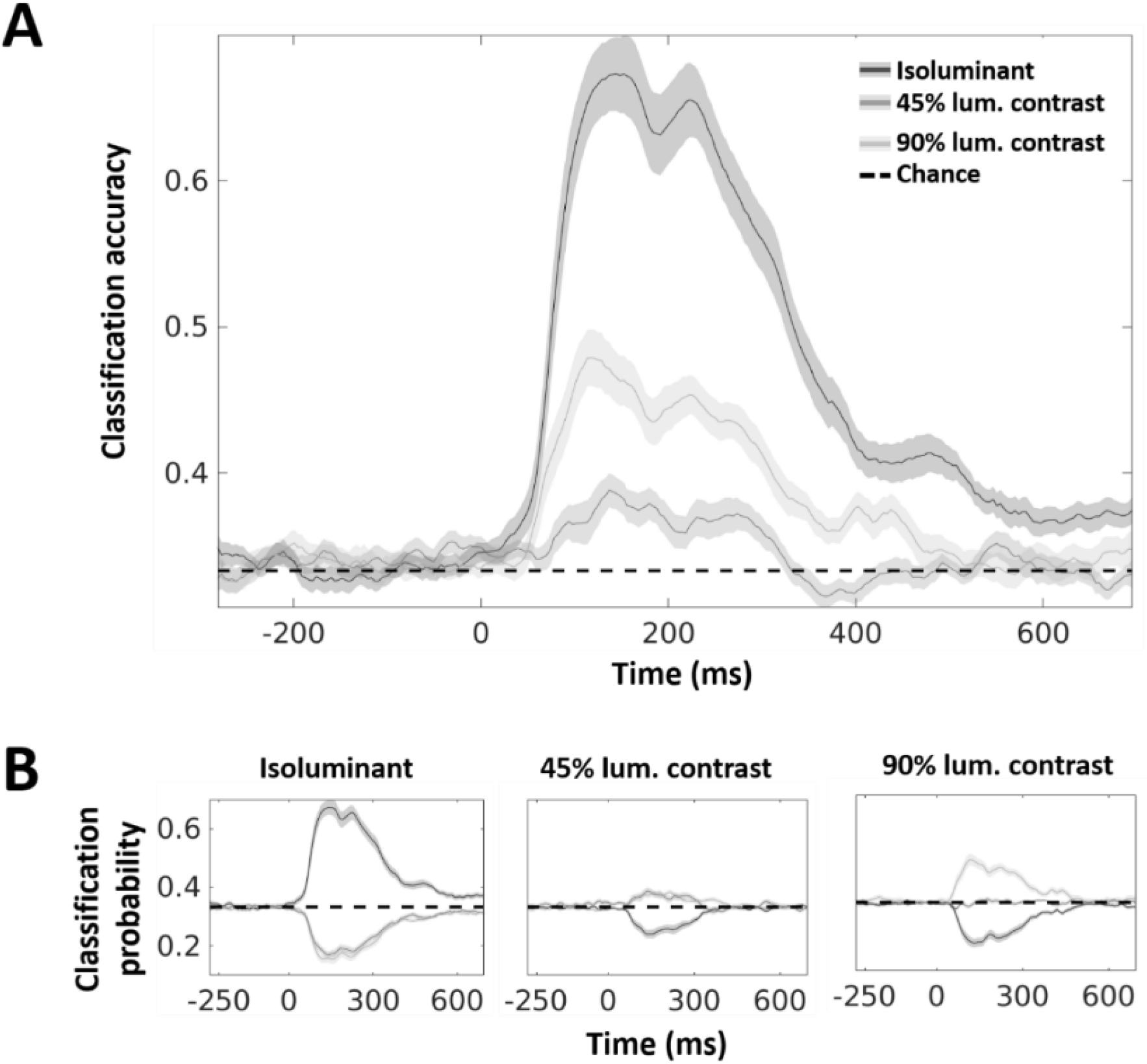
Luminance decoding from EEG signals. A. Mean classification accuracy. This panel shows the performance of the model in correctly identifying the luminance contrast of the stimuli (model accuracy). Each line shows the accuracy for one condition, with dark grey coding for the isoluminant condition, medium grey coding for 45% luminance contrast, and light grey coding for 90% luminance contrast (coding of luminance contrast using lightness is used throughout the article). The shaded area around the lines shows ±1 standard error of the mean. Chance performance is shown by the dashed line. B. Misclassification probabilities. Each subpanel shows one row of the confusion matrix analogous to Figure 2C. The left panel shows classification probabilities for the three luminance conditions when isoluminant stimulus is presented to the classifier. Similarly, the middle and right panels show prediction probabilities when 45% and 90% luminance contrast inputs are presented to the classifier.

Stimuli with 45% luminance contrast have an above-chance probability of being misclassified as 90% luminance contrast (and vice-versa). However, this effect is not strictly symmetric, with 90% luminance contrast being easier to detect compared to the 45% contrast. Thus, under non-isoluminant conditions, not only are the hue-driven patterns difficult to detect, but they seem to be progressively overridden by luminance-contrast-driven patterns. To ensure that this effect was driven by luminance, and not by the chromatic content of the stimuli, we set up separate models for each hue, and were able to confirm that the effect was indeed independent of the chromatic content of the stimulus. For each hue, the isoluminant stimuli were robustly classified (Supplementary **Figure S2**, leftmost column), while the non-isoluminant conditions produced similar but asymmetric prediction scores (Supplementary **Figure S2**, middle and right columns).

### Interim Discussion

Our findings are in line with Sutterer et al (2021) who recently reported that both colour and luminance content can be successfully decoded from EEG signals. Hermann et al (2021) investigated decoding of hue or luminance polarity from MEG signals and found that generalising luminance polarity across hue works better than generalising hue across polarity. This is consistent with our own findings that decoding of hue is strongly affected by the addition of luminance contrast. Unlike these studies, where only stimuli that combine colour and luminance contrast were used, we also included stimuli that were isoluminant with the background. We found that decoding of hue from such nominally isoluminant stimuli is much more efficient. While it appears that decoding was superior for unique compared to intermediate hues, Hermann et al (2021) also report higher decoding efficiency for red and green compared to orange and blue, although such an asymmetry is not present in the decoding study by Hajonides et al. (2021). Hermann and colleagues suggest that poorer decoding for orange and blue may be due to their alignment with the daylight locus, causing a less consistent signal in the presence of luminance. To disambiguate if unique or intermediate hue status drives a more robust neural signal irrespective of daylight locus alignment, it would be necessary to use a unique hue that is also more aligned with the daylight locus, such as yellow or blue. Thus, in our next experiment, we decided to replace red with yellow, which would allow us to maintain the same proximity structure (red/orange, green/turquoise) but eliminate the potential daylight locus confound.

Finally, the stimulus set in Experiment 1 was designed to investigate whether unique hues have more robust EEG representations. To achieve this, we chose unique and non-unique hues that were maximally distant in a perceptual space – red and green, orange and turquoise (see details of the stimulus set in Methods). As already reported by Rosenthal et al (2021) and Hermann et al (2021), inter-hue differences in decoding efficiency manifest even between such evenly spaced colours. To better understand the non-uniformity of this neurometric colour space, in our next experiment we aimed to investigate the structure of the decoding manifold by introducing proximal neighbours, clockwise and counterclockwise to each hue. Decoding colours in such small and large neighbourhoods allows us to understand how perceptual notions of hue-difference map to the EEG-derived neurometric space.

### Experiment 2: Decoding over small and large perceptual hue differences

In Experiment 1 we showed superior decoding performance for unique hues compared to intermediate hues, suggesting a robust neural representation for the former. In Experiment 2, this hypothesis was further critically tested by using small and large hue differences. Our aim was to re-examine decoding of nominally isoluminant unique and intermediate hues with a slightly modified hue set (see Interim Discussion above) and to extend it by decoding local clusters of stimuli around each of these hues. First, we measured individual settings for unique (yellow and green) and non-unique (orange and turquoise) hues for each observer. Next, we made EEG measurements in a task analogous to Experiment 1 using, for each observer, a stimulus set consisting of their subjective settings for the four hues (denoted as the = configuration), and two sets of stimuli generated by rotating the subjective settings by ±10° in CIELAB colour space (denoted as the + and – configurations respectively) – leading to a total of 12 stimuli (4 hue-clusters and 3 rotational-configurations, see Supplementary **Figure S5D**). The individual hue settings were as follows (means and SEs): yellow 101° ± 2°, orange 61° ± 3°, green 153° ± 3° and turquoise 198° ± 3°.

In the shape discrimination task, grand mean accuracy was 96% ± 1 % SE (see Supplementary **Figure S5A**) and reaction times were 706 ± 61 ms (See Supplementary **Figure S5B**). Response-time data was analysed with a 4×3 repeated measures ANOVA (4 hues vs. 3 rotational configurations, i.e., −, + and = sets), which yielded a significant main effect of hue (F (1.77, 26.5) = 5.25, p=.01, ηp2 = 0.26) and an interaction with the rotational configuration (F (2.16, 32.49) = 5.08, p =.01, ηp2 = 0.25) while the effect of the rotation itself was not significant (F (1.79, 26.99) = 0.72, p = 0.48, ηp2 = 0.05). The interaction was deconstructed by separate repeated measures ANOVAs at each hue: for yellow, there was a significant effect of rotation (F(1.36,20.48) = 6.23, p =.01, ηp2 = 0.29) driven by slower RTs for the individual hue setting vs. 10° clockwise setting (p = 0.006). For green, there was also a significant effect (F(1.58,23.74) = 6.76, p =.007, ηp2 = 0.31) driven by faster RTs for the individual hue setting vs. 10° clockwise setting (p = .04) as well as vs. 10° counterclockwise setting (p = .005); no differences were found for orange (p=.22) and for turquoise (p=.11). Taken together, we can see that only for unique hues (yellow and green) the responses to individual hue settings (= configuration) seem to be different from responses to ±10° rotated hues (i.e., – and + configurations). However, the direction of the effect was opposite for the two hues – while participant responded slower to their individual yellow setting, they responded faster to their individual green setting.

For the Categorical Rating task, the average ratings and their SEs were as follows: individual yellow 5.62 ± 0.6; −15° yellow 5.75 ± 0.57; +15° yellow 2.56 ± 0.35; individual green 6.93 ± 0.26; −15° green 7.93 ± 0.17; +15° green 3.87 ± 0.35; individual orange 6.37 ± 0.36; −15° orange 3.93 ± 0.48; +15° orange 7.25 ± 0.48; individual turquoise 5.68 ± 0.53; −15° turquoise 7.43 ± 0.53; +15° turquoise 3.18 ± 0.5 (See Supplementary **Figure S5C**).

#### Decoding over large hue differences is predicted by hue angles

For each observer, we trained tECOC models over all stimuli: the four hue settings (= group), and the eight stimuli generated by ±10° rotations of each of these settings (+ and - groups respectively). Using the classification results, we generated a time-series of dissimilarity matrices (see Methods for details) and found that the stimulus representations were highly dissimilar in a 100-400 ms window after stimulus onset (**Figure 6A**). Similarly, we also calculated a perceptual dissimilarity measure by using differences in hue angles of the stimuli in CIELAB space. As expected, perceptual dissimilarity increases as one moves away from a given reference stimulus (**Figure 6B**). Using rank-correlation analysis, we found a significant (*p* < 0.001) increase in Kendall’s tau statistic in a 100-400 ms range post-stimulus (**Figure 6D**), suggesting that perceptual distances are indeed correlated with decoding output. This was also reflected in stable mean and peak dissimilarities during the period of significant correlation (**Figure 6C**).

**Figure 6:**
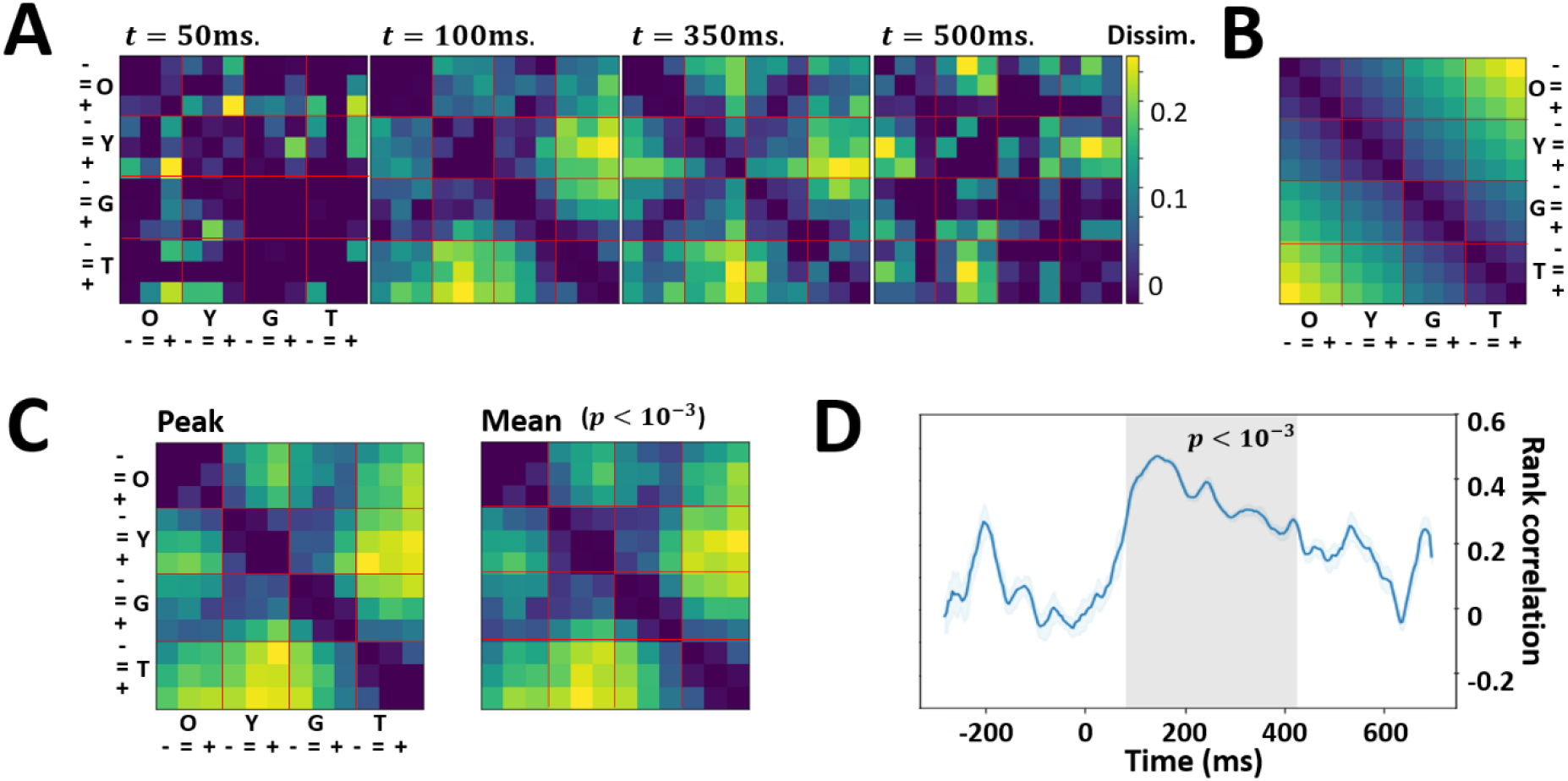
Isomorphism between representational and perceptual spaces for large distances. **A.** Dissimilarity in classifier outputs. tECOC models were trained to classify twelve colours from their EEG responses. The colours sampled four clusters along the hue circle corresponding to orange (O), yellow (Y), green (G), and turquoise (T), with each cluster consisting of settings made by the observer in a psychophysical experiment (=), and colours sampled 10° clockwise (−) and anti-clockwise (+) with respect to each setting. Each panel shows a dissimilarity matrix derived from classifier output. The panels show the dissimilarity 50, 100, 350, and 500 ms after stimulus onset. **B.** Dissimilarity in perceptual space. Hue angles of the 12 stimuli (same as panel **A**) were used to estimate dissimilarity in the perceptual CIELAB space. **C.** Peak and Mean Dissimilarity. The left panel (labelled Peak) shows the dissimilarity is classifier output at the time-point where rank-correlation (panel **D**) with perceptual dissimilarity peaks, and the right panel (labelled Mean) shows the average dissimilarity over the period where the correlation is statistically significant. **D**. Representational similarity. Rank-correlation (left panel) between perceptual and classification dissimilarities using Kendall’s tau statistic. Grey background indicates statistical significance (*p* < 0.001), and the statistic peaks at *t* = 145 ms.

#### Local distortions in hue decoding

Next, we posed the question: is the perceptual robustness of unique hues reflected in the structure of the decoding space around their respective representations? To answer this question, we trained 4 tECOC models – one in the neighbourhood of each of the four individually measured hues. Each model was trained to classify EEG signals into one of three labels: subjective setting (=), stimuli 10° clockwise (+) to the subjective settings, and stimuli 10° counter-clockwise (−) from the subjective settings. In **Figure 7A** we show the results for the four models, one model per row. Each subpanel is a row in the corresponding confusion matrix, with the test stimulus indicated on top – for instance, the first panel shows the predictions of the model trained in the yellow neighbourhood, when stimuli 10° counter-clockwise from subjective yellow settings are presented to it.

**Figure 7:**
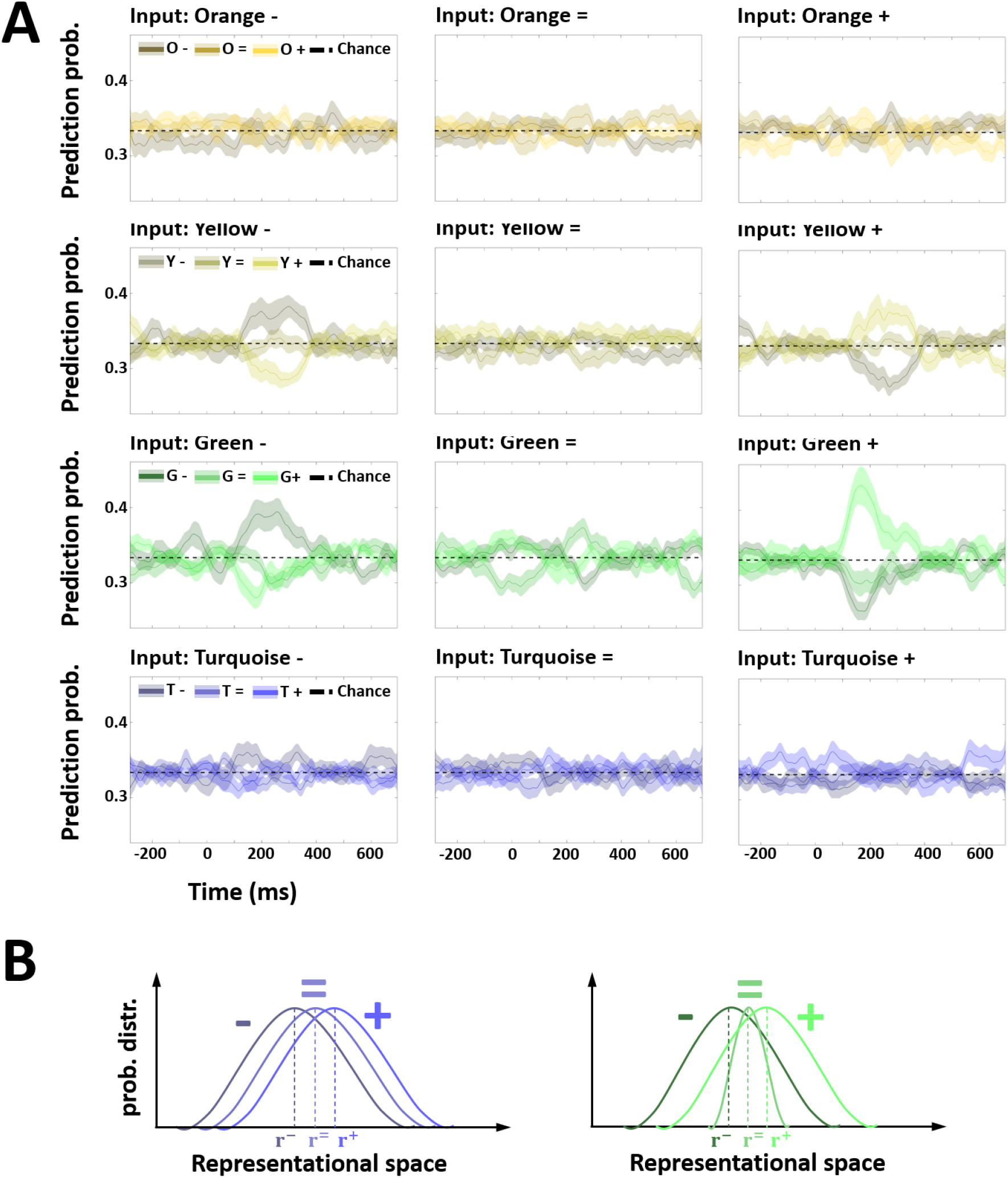
Local distortions in representational space. **A.** Decoding within colour clusters. tECOC models were trained on clusters around the subjective settings for unique (yellow: Y, green: G) and non-unique (Orange: O, Turquoise: T) hues. Each cluster consisted of three groups: subjective observer settings (=), and two groups derived from 10° clockwise (+) and counter-clockwise (−) rotations of the subjective settings in CIELAB space. Each row shows a model trained on a different hue (top row: yellow, second row: green, etc.), with subpanels showing rows of the corresponding confusion matrices. The input stimulus for each row of the confusion matrix is labelled on top. **B.** Effect of variability in the internal neural representation ***r*** on decoding. The panel on the left shows a configuration of the distributions for the three groups (−, = and +) which can lead to poor decoding scores such as those observed for non-unique hues. The distributions overlap, and the distance between the distributions is too low to allow for proper discrimination using a linear boundary. The panel on the right shows how a relative decrease in the variability of one group (subjective settings for unique hues, =) can lead to better decoding for the other two (− and +).

We found that the three groups (subjective settings, and the ±10° rotations) cannot be decoded in non-unique hues (**Figure 7A**, first and fourth rows). However, for unique hues (**Figure 7A**, second and third row), the rotated groups (first and third columns) can be decoded, while the subjective settings (second column) cannot. This result suggests that EEG representations of unique hues may have lower variability compared to non-unique hues. Such fluctuations in local variability in the representational space can create distortions in the decoding measure, allowing for better decoding of the flanking distributions (one such scenario is illustrated in **Figure 7B**). Note that in the perceptually uniform CIELAB space the three groups, by design, had equivalent relative distributions (− and + were simply mean-shifted copies of =).

## Discussion

Our first finding is that - under isoluminant conditions - EEG responses to unique hues show more distinct patterns compared to non-unique hues, and that these patterns are stable during both passive viewing (**Figure 2**) and active task-engagement (**Figure 3**). We can also reach certain conclusions about the underlying neural processes from the time-course of decoding performance. A 100-300 ms decoding window is consistent with the idea that the performance of the model could be driven by both perceptual and post-perceptual contributions (Forder et al., 2017b). This is supported by the fact that the decoding performance steadily rises before peaking between 150-200 ms after stimulus onset, a time-window where EEG signals begin reflecting post-visual evaluative processing (VanRullen and Thorpe, 2001), including colour categorisation (Fonteneau and Davidoff, 2007). The chromatic visual evoked potential (cVEP), which reflects the activation of colour sensitive neurons in early visual cortices, also remains maximal in the same time window (Nunez et al., 2018). However, a high-level interpretation of the decoding on the basis of the categorical status of the stimulus colours is unlikely. Categorical representativeness ratings do not follow the pattern observed in the classifier performance (see Supplementary **Figure S4C**), and seem to rather reflect the relation between the colour sample and the focal colour. The most parsimonious explanation for the pivots in colour space that drive asymmetries in decoding around unique hue locations would be that they correspond to hue locations that are associated with a more robust neural representation, thus making it more easily decodable from less robustly represented hues.

Secondly, classification performance for the decoding of hues is diminished when luminance contrast was added (**Figure 4**). This was not entirely unexpected since luminance contrast is known to have a strong effect on EEG responses, once luminance contrast is sufficiently strong (Rabin et al., 1994). At the same time, we found that all luminance conditions **Figure 5** can be decoded to above-chance levels within the same 100-300 ms window. Thus, under non-isoluminant conditions, not only are the hue-driven patterns more difficult to detect, but they may also be at least partly overridden or replaced by luminance-contrast or joint-colour-and-luminance-contrast-driven activity. Our findings are consistent with the idea that hue is most likely to be encoded by neural populations which also encode luminance. The fact that purely chromatic-tuned cells in the visual cortex are known to be in a minority compared to luminance-tuned or luminance-chromaticity tuned cells (Lennie et al., 1990; Johnson et al., 2001) may partly explain why luminance signals tend to override chromatic information in EEG recordings. In V1-V3, the neurons are tuned to many intermediate directions, both in terms of hue and luminance contrast (for a review, see Gegenfurtner and Kiper, 2003). In higher-level areas of the extra-striate cortex, colour representations become organised in ways that resemble perceptual colour spaces (Brouwer and Heeger, 2009, 2013). Thus, the decoding in our study is likely to reflect cumulative effects that build up across these areas. Even though we find more robust responses for the two unique hues (red and green) compared to the two non-unique hues (orange and turquoise), decoding is still possible for non-unique hues, implying that there are indeed multiple hue representations that are being encoded by the brain (see, e.g., Brouwer and Heeger, 2009; Parkes et al., 2009; Zaidi and Conway, 2019).

Thirdly, we show in Experiment 2, that the geometric structure of this representational space can be explored by carefully designed experiments. Our results demonstrate that while large distances in the neural representational space are indeed correlated with perceptual hue differences (**Figure 6**), there are local anisotropies associated with unique hues (**Figure 7**) which are likely to represent local changes in signal variability. Such tunings could reflect properties of our environment such as the statistical regularities in the reflectance spectra of naturally occurring surfaces (Philipona and O’Regan, 2006). Perhaps this is the reason why the neural reality of perceptual red-green and blue-yellow hue-opponent mechanisms has proven to be so elusive – it is not a fundamental mechanism hard-wired into the neural circuitry, but a statistical peak in the tuning of neural populations which multiplex both colour and luminance information. Its identification is therefore complicated by the fact that neural populations jointly coding for chromaticity and luminance are likely to show higher responsiveness to the presence of luminance contrast (Johnson et al., 2001), making hue-specific signals much harder to detect.

A growing number of studies investigating population activity analyse EEG and MEG topographical data by interrogating trajectories in activation manifolds. Our results suggest that the structure of such manifolds can be highly anisotropic, and that these anisotropies can reflect perceptual measurables. In the case of hue perception, it is likely that the local structure of this space is reflected in quasi-invariants such as the so-called unique hue percepts. Now that neurometric mapping of hue spaces has been established by numerous studies (Hajonides et al., 2021; Hermann et al., 2021; Rosenthal et al., 2021), this study marks a first hypothesis-based exploration of these maps and shows that unique hues represent local anisotropies in cortical hue-representations.

### Open practices

The decoding scripts have been packaged as the tECOC toolbox, which has been made available as a public git repository here. The EEG and behavioural data from both experiments will be shared on the Open Science Framework website.

## Acknowledgments

TC was partly supported by grant FRM: SPF20170938752 (Fondation pour la Recherche Médicale). JM was partly supported by grants BB/H019731/1 and BB/R009287/1 from the Biotechnology and Biological Sciences Research Council (BBSRC).

## Supplementary Material

### S1: Model robustness

**Figure S1.**
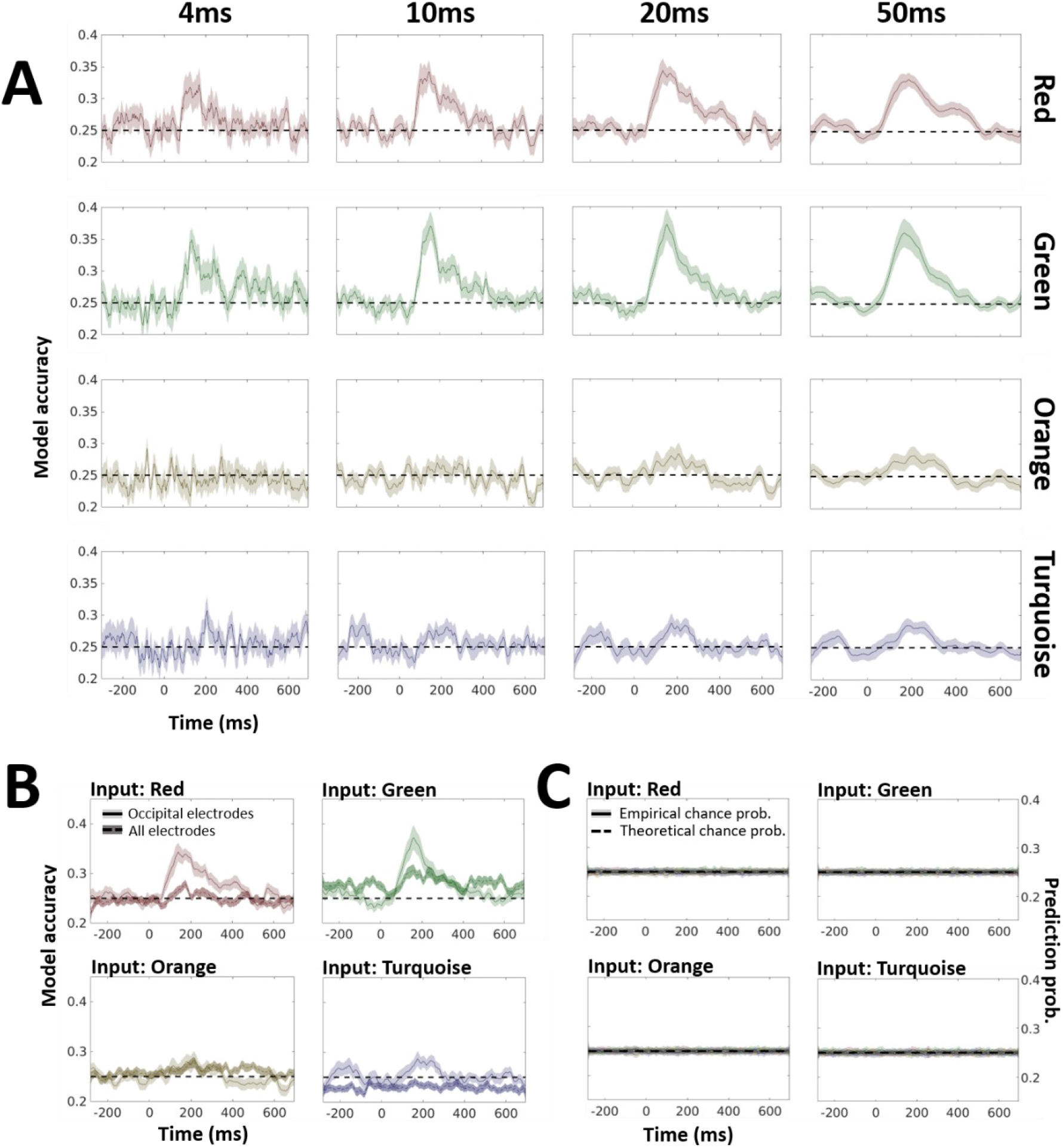
Model robustness. **A.** Effect of time-window on model performance. Isoluminant stimuli were used to train models using time-window lengths (see Methods for details) of 4ms, 10ms, 20ms and 50ms. Each subpanel shows the performance for one such model (columns, labelled on top) in correctly identifying the label corresponding to the input EEG signal (rows, labelled to the right). The mean accuracy is shown as a solid line, while the shaded envelope indicates standard error of the mean. A dashed line is used to show the theoretical chance level. **B.** Decoding using all electrodes. The classification of isoluminant stimuli was performed using all 64 electrodes in a 20ms window. The results (dotted lines with darker envelopes) are shown along with those obtained by using only the occipital electrodes (solid lines with lighter envelopes). The lines indicate mean performance while the shaded envelopes indicate standard error of the mean. The theoretical chance level is indicated by dashed horizontal lines. **C.** Comparison of theoretical and empirical chance performance. To calculate the empirical chance performance, we repeated the experiment shown in Figure 2 using shuffled labels for training the model. This ‘permuted’ model was then tested using the same sequence of stimuli used for testing the unshuffled trained model. Each subpanel in the figure shows the prediction probabilities for one particular input hue (same analysis as Figure 2C).

Here we demonstrate that our findings are robust to key parameters. First, we show that the time-window is not a critical parameter for our analysis (panel **A**). The classifier performance does not change qualitatively with an increase/decrease of the time window length. However, we do observe increasing noise in the model performance as the window-length approaches the sampling frequency. Second, we show the classification accuracy for a model which was trained on isoluminant stimuli using all 64 channels of the EEG signal (panel **B**). As expected, the results show similar trends to those found in **Figure 2**. The decrease in performance could be due to the added noise in the hue-related signal from non-occipital sites. Finally, we show the results from a permutation analysis (the model is trained using a shuffled set of labels and tested with correctly labelled data) showing that empirical chance-performance is very close to the assumption that all labels are equally likely.

### S2. Robustness of luminance decoding across hues

In **Figure 5** we show the results of a simulation where EEG signals were used to decode the luminance of the stimulus. Here, we show results from additional simulations which demonstrate that this was not driven by any particular hue. Four models were trained to classify the luminance, one for each of the four hues (rows). For each of the four models, the isoluminant stimuli (left column) were robustly identified, while the non-isoluminant conditions (middle and right columns) were most likely to be confused with one another. We also observe the asymmetry between the 90% and 45% luminance-contrast conditions where the former is easier to detect than the latter.

**Figure S2.**
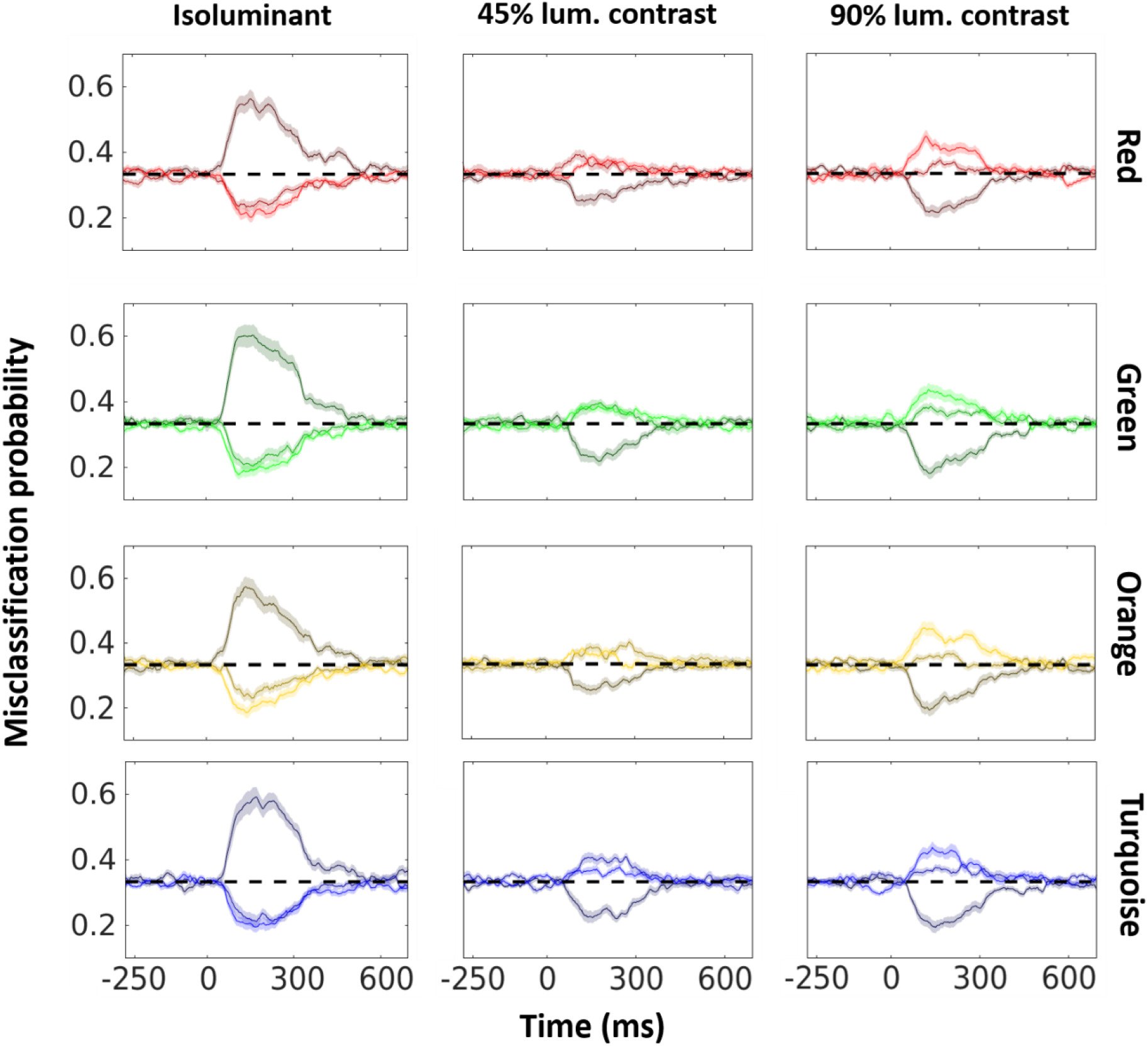
Luminance classification is robust across hues. ERP responses to individual hues were used to train hue-specific models for luminance contrast classification. Each row shows the results for one of the four models. In each model (row), the input luminance contrast is denoted by the column (left: isoluminant, middle: 45%, and right: 90%), and lines in each panel denote the probability of the predicted label (coded by the lightness of the colour – lowest lightness: isoluminant, medium lightness: 45%, and high lightness: 90%). The shaded area around the mean represents standard error of the mean. Dashed lines show the theoretical chance performance for the models.

### S3: Stimulus coordinates

Here, we show the coordinates of the stimuli for both experiments. Experiment 1 used nominal unique hues based on a large dataset (*N* = 185) of settings, and non-unique hues which were equidistant from their respective closest unique hues. Experiment 2 used subjective settings for both unique and non-unique hues, measured in separate sessions before the EEG recordings. See Methods for details.

**Figure S3.**
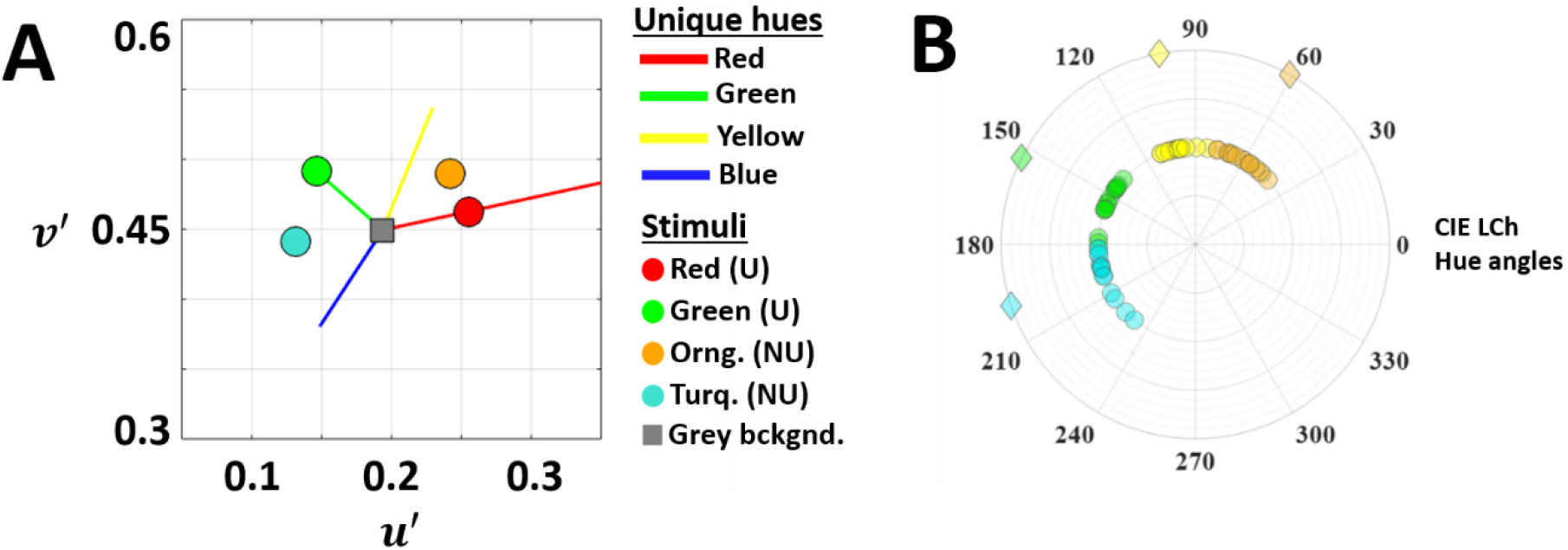
Stimulus coordinates. **A.** Experiment 1. Mean observer settings from a large dataset (*N* = 185) of uniuqe hues were used to select the unique red (UR) and green (UG) stimuli. The non-unique hue stimuli were chosen such that orange was equidistant (in terms of hue angle) from UR and unique yellow (UY), and turqoise was mid-way between UG and unique blue (UB). All stimuli had the same saturation level (set equal to the maximum possible saturation for UG within the monitor gamut). The axes are the u’ and v’ coordinates in the perceptually uniform CIE 1976 UCS space. The mean hue angles for each unique hue from the dataset are shown as solid lines. The length of these lines indicates the limits of the monitor gamut. The stimuli (unique: U, non-unique: NU) are shown as coloured dots, while the background is shown as a grey square. **B.** Experiment 2. Polar plot showing participants’ individual settings for unique (yellow and green) and intermediate hues (orange and turquoise) in CIE LCh colour space. Circles represent participants’ individual settings and diamonds represent the mean of those settings

### S4: Behavioural measurements in Experiment 1

During the experiment we also measured accuracy and reaction times on the shape change task, while at the end of the experiment we captured a category representativeness rating for each colour (see *Methods* and *Results* for details). Here we show that these measures do not distinguish between unique and non-unique hues. Furthermore, representativeness ratings provided by the participants indicate that across all conditions, the turquoise and green stimuli were judged to be the most representative of their colour name (panel **C**). Taken together, we see that behavioural measurements do not correlate with whether a given stimulus was unique or non-unique.

**Figure S4.**
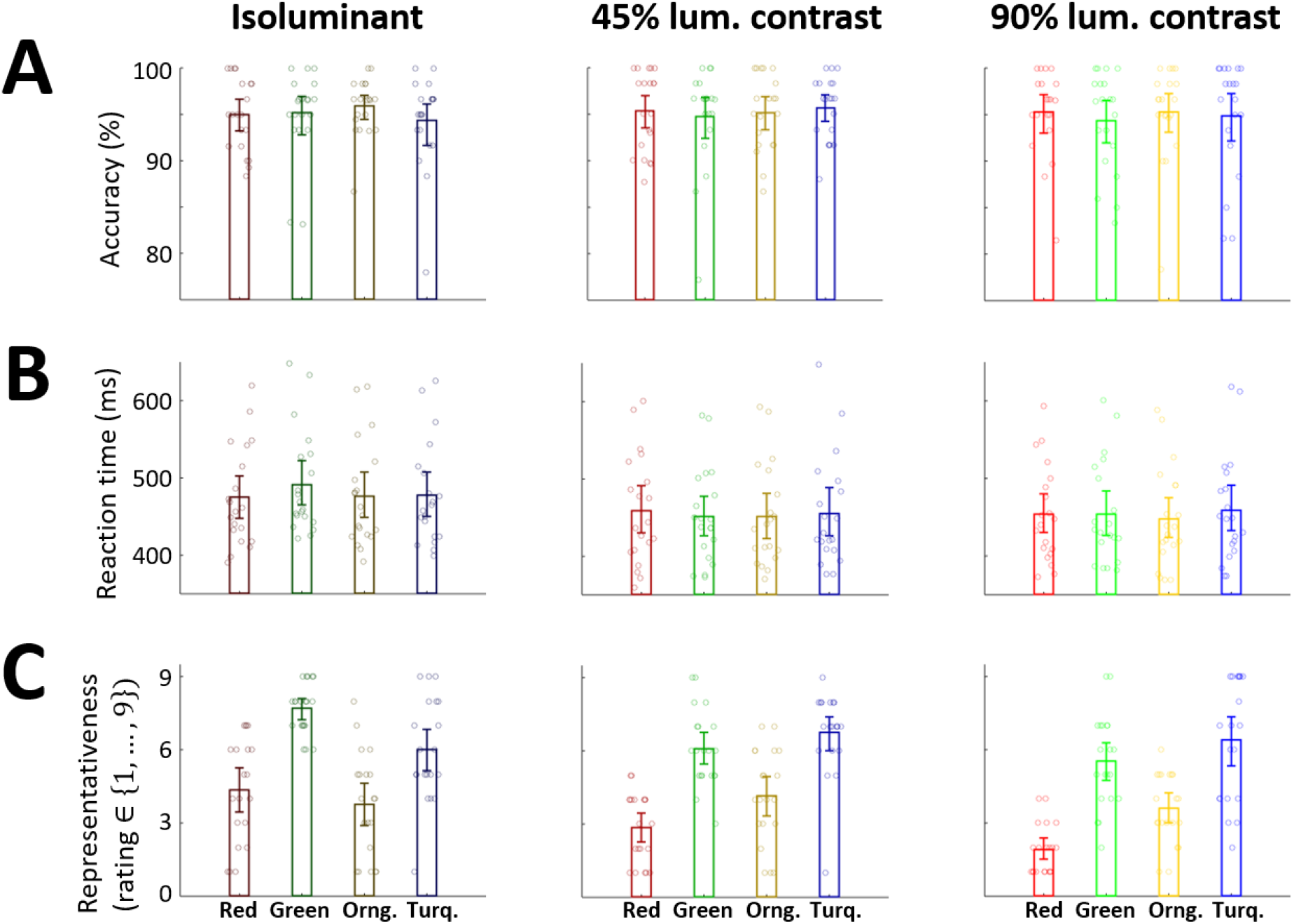
Behavioural results do not reflect unique/non-unique hue status. **A.** Task accurarcy. During the shape-change segment, the participants were asked to report whether a circular stimulus changed to a square or a diamond (see *Methods*). The participant accuracy is reported for reach condition. **B.** Reaction time. During this task, the reaction times were also recorded. **C.** Representativeness ratings. The participants also provided representativeness ratings for the stimuli. E.g., if a red stimulus was being presented, we asked the participant to rate how this stimulus represented the category ‘red’. Ratings were reported on a Lickert scale going from 1 to 9. In all panels, the height of the bar represents the mean, the error bars represent bootstrapped 95% confidence intervals (1000 samples were drawn), and circles show the raw data (20 participants).

### S5: Behavioural measurements in Experiment 2

**Figure S5.**
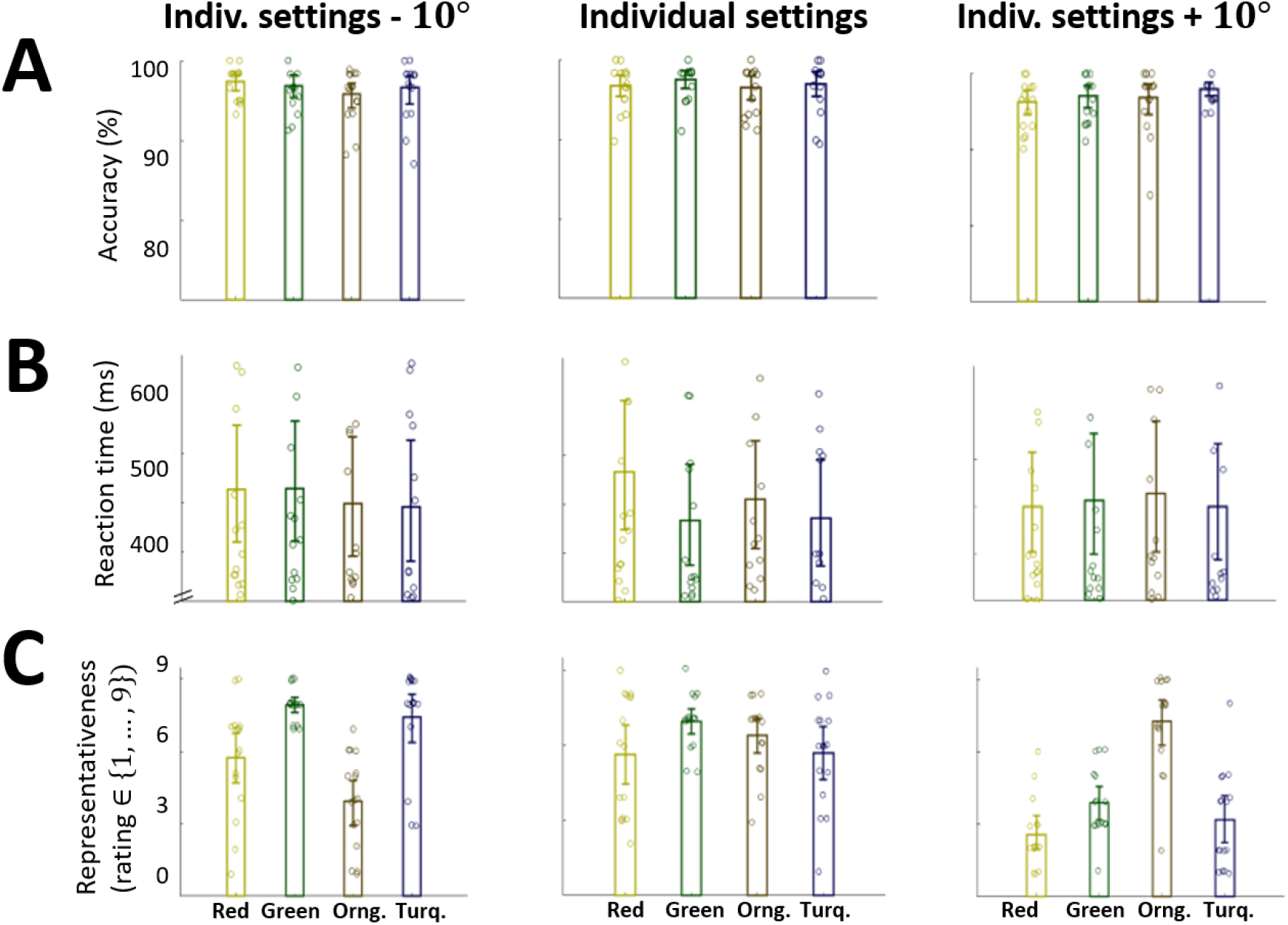
Behavioural results from Experiment 2. **A.** Task accurarcy in the shape discrimination task. **B.** Reaction time recorded during this task. **C.** Categorical representativeness ratings (16 participants). The panels use the same conventions as **Figure S4A**, **B** and **C** respectively.

### V1: Time-course of the confusion matrix

Here, we show how the confusion matrix for isoluminant stimuli changes over time. Between 100-300ms, unique red and green show increased prediction probabilities, while the non-unique hues do not. Furthermore, for any presented stimulus, positive deviations from chance are only seen for the correct label and the proximal hue but not for non-proximal hues (e.g. for a red stimulus, the model predicts the labels red or orange, but never green or turquoise).

**Link**: https://www.dropbox.com/s/b6fzc731fbmksu5/Classification_UH_Isoluminant.avi?dl=0

**Video V1:** The label of the presented stimulus is shown on the x-axis, while the y-axis shows the label predicted by the model (thus, correct predictions lie on the diagonal going from bottom-left to top-right). The colour of a given square at time *t* represents the predicted label at this instant, and the intensity of the colour shows the deviation of the prediction probability from the theoretical chance-level. For clarity of presentation, any negative deviations from chance have been clipped to zero. The time elapsed from stimulus onset is shown on top.

